# Chemical-induced degradation of PreS2 mutant surface antigen reverses HBV-mediated hepatocarcinogenesis

**DOI:** 10.1101/2022.01.24.477454

**Authors:** Joey Yi Yang, Yi-Hsuan Wu, Max Yu-Chen Pan, Yu-Ting Chiou, Richard Kuan-Lin Lee, Tian-Neng Li, Lily Hui-Ching Wang

## Abstract

Naturally evolved immune-escape PreS2 mutant is an oncogenic caveat of liver cirrhosis and hepatocellular carcinoma (HCC) during chronic hepatitis B virus (HBV) infection. Notably, PreS2 mutants is prevalent in above 50% of patients with HCC. Intrahepatic expression of PreS2 mutant large surface antigen (PreS2-LHBS) induces endoplasmic reticulum stress, mitochondria dysfunction, cytokinesis failure and subsequent chromosome hyperploidy. In this study, we ask if long-term inhibition of PreS2-LHBS may act to reverse HBV-mediated hepatocarcinogenesis. We set up a stability reporter platform and identified ABT199 as an inhibitor of PreS2-LHBS from a library of 1068 FDA-approved drugs. Treatment of ABT199 induced PreS2-LHBS degradation without affecting the general cell viability, as shown in hepatoma and immortalized hepatocyte cell lines. We found that ABT199 induced the recruitment of PreS2-LHBS to ring-shaped structures in close proximity to lysosomal marker Lamp1 and multivesicular body marker Rab7. Simultaneously, inhibitions of lysosomal degradation or microautophagy restored the expression of PreS2-LHBS. Specifically, a 24-hr treatment of ABT199 reduced DNA damages and cytokinesis failure induced by PreS2-LHBS. Persistent treatment of ABT199 for 3 weeks reversed chromosome hyperploidy in PreS2-LHBS cells and suppressed anchorage-independent growth of HBV-positive hepatoma cell line. Together, we showed that ABT199 provoked selective degradation of PreS2-LHBS through the induction of microautophagy, and a long-term treatment of ABT199 reversed oncogenic mechanisms induced by HBV. Our results indicate that long-term degradation of PreS2-LHBS may serve as a novel therapeutic strategy to constrain HBV-mediated hepatocarcinogenesis.

**Graphical abstract:** 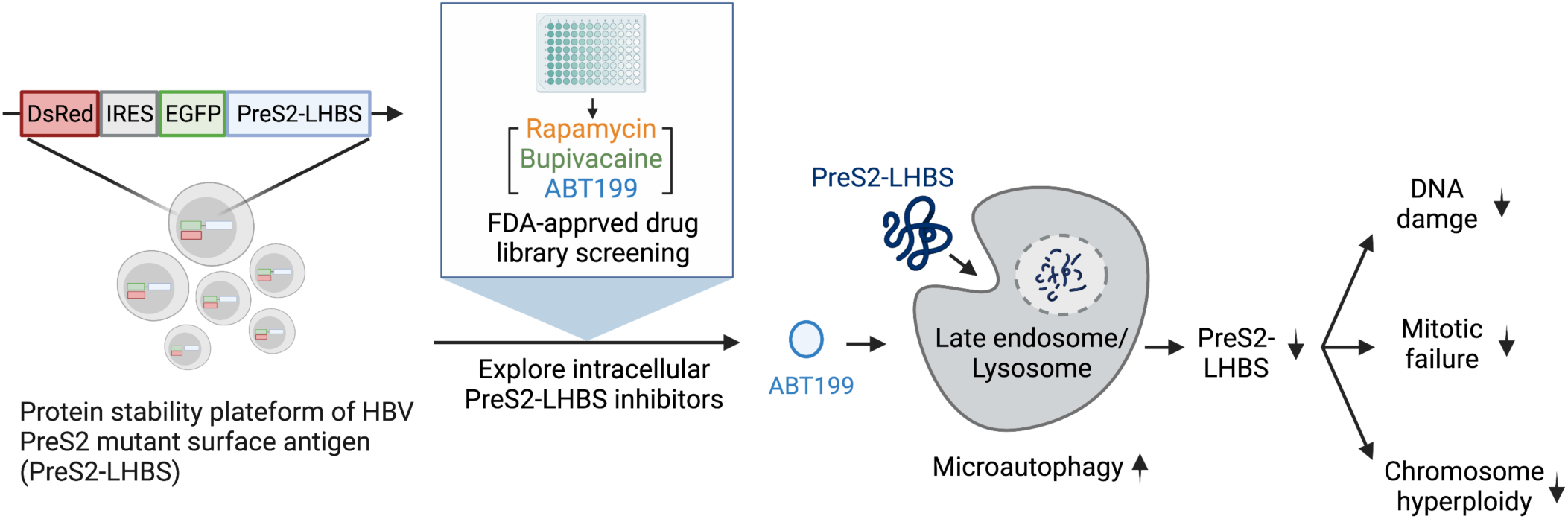

**Author summary:** HBV PreS2 mutant large surface antigen (PreS2-LHBS) is an oncoprotein that induces liver cirrhosis and liver cancer. This study identified ABT199 as a potential inhibitor of PreS2-LHBS. We found that ABT199 could trigger the degradation of PreS2-LHBS through the induction of microautophagy. Moreover, a long-term treatment of ABT199 significantly reversed LHBS-induced oncogenic events including DNA damage, mitotic failure, chromosome hyperploidy, and anchorage-independent growth. This study not only identifies a specific inhibitor of PreS2-LHBS but also highlights a plausible strategy to constrain HBV-mediated hepatocarcinogenesis through targeting the degradation of PreS2-LHBS.

## Introduction

Chronic hepatitis B virus (HBV) infection is the most prevalent cause of liver diseases—3.5% of people were chronically infected worldwide; 20-30% of patients progress to liver cancer (hepatocellular carcinoma, HCC) and liver cirrhosis; 25% of patients eventually died in the diseases attributed to chronic HBV infection [1,2]. Although a prophylactic vaccine and antiviral drug exist, no cure at this stage [1,2].

HBV S gene encodes three segments (PreS1, PreS2, S) that translate three proteins differing in length but sharing the same C terminal: large (LHBS), middle (MHBS), and small surface antigen (SHBS). All three surface antigens together constitute HBV surface antigens (HBsAg) in serology. Surface antigen plays a role in mediating disease progression in CHB patients. A high level of HBsAg has been found to suppress CD8 positive T cell function, leading to T cell exhaustion and incomplete virus clearance [3,4]. In addition, viral surface antigen has been identified as an oncoprotein associated with HCC and cirrhosis progression. Transgenic mice overexpressing surface antigen have been found to develop HCC [5]. In several studies, patients with a high HBsAg level (>1000 IU/mL) showed increased risk of HCC even in patients with low HBV DNA level (<2000 IU/mL) [6–8]. We recently also showed that LHBS might overwrite hepatocyte G2/M checkpoint, increase DNA damage, induce cytokinesis failure, and consequently provoke hepatocyte hyperploidy [9,10].

During the chronic hepatitis B (CHB) progression, surface antigen accumulates in ER, leading to flat and hazy cell morphology, called ground-glass hepatocyte (GGH). According to immunohistochemistry staining of surface antigen, GGH can be classified as type I and type II GGH. Type I GGH displays a homogenous cytoplasmic distribution of surface antigen; type II GGH displays a marginal distribution of surface antigens. In CHB patients, the prevalence of type II GGH increases upon disease progression. This advanced stage of GGH usually bears a mutant LHBS with PreS2 region deletion (PreS2-LHBS, predominate deletion in 2-55 or 4-57) [11]. Studies have shown that type II GGH carrying PreS2-LHBS has a significant association with HCC, cirrhosis, and the decreased survival rate [12,13]. In addition, compared to wild-type LHBS, PreS2-LHBS shows higher ER retention that induces a substantial ER stress, leading to calcium accumulation, mitochondrial dysfunction, and genomic instability [14,15]. Notably, the PreS2 deletion region corresponded to the epitope of CD8 T cell, suggesting the immune-escape attribute of this mutant [11]. Furthermore, transgenic mice expressing PreS2-LHBS are sufficient to develop liver dysplasia and HCC [16]. Notice that the increasing prevalence of integrated HBV genome, a driving force of HCC, particularly supports the consistent expression of intracellular surface antigen [17]. These studies indicate that intrahepatic PreS2-LHBS is not only a histological hallmark but a caveat of disease progression to liver cirrhosis and HCC. Thus, targeting PreS2-LHBS is a therapeutic option to constrain HBV-induced hepatocarcinogenesis.

Therapeutic options to eliminate HBV are not available at this stage. The ideal treatment endpoint, called “functional cure”, is determined by the complete loss of HBsAg in the circulation. According to the AASLD hepatitis B guideline in 2018, approximately only 1∼ 8% of HBeAg-positive patients achieved HBsAg loss with current treatment interferon and nucleotide analog (NA). The chance of HBsAg loss was further down to below 1% in HBeAg-negative patients [18]. Note that even though NA significantly reduced the viral marker in serum, the expression of intrahepatic virus HBsAg was not be affected, especially the type II GGH carrying viral preS2 mutant, which was linked to decreased overall survival of HCC [19]. Coordinately, PreS2-LHBS was detected in above 60% of HCC patients [11]. Accordingly, intrahepatic PreS2-LHBS is a risk factor for hepatocarcinogenesis, and yet no conventional therapeutic can eliminate such oncogenic caveat in the liver at this stage.

As a result, there is an unmet medical need for the reduction of intrahepatic PreS2-LHBS in CHB patients. We ask if long-term inhibition of PreS2-LHBS may reverse the existing oncogenic effect. Therefore, we set up a protein stability platform to screen for chemical inhibitors that induce the degradation of intracellular PreS2-LHBS. From an FDA-approved drug library, we identified ABT199 as a candidate inhibitor of PreS2-LHBS. We showed that long-term degradation of PreS2-LHBS was achieved by treating with ABT199, which subsequently reversed oncogenic effects induced by PreS2-LHBS. These results indicate that the long-term degradation of intrahepatic PreS2-LHBS may serve as a novel therapeutic approach to constrain the development of HBV-mediated HCC.

## Result

### Repurposing FDA-approved drug to target PreS2-LHBS through protein degradation

To target PreS2-LHBS, we decided to explore the existing chemicals that reduce PreS2-LHBS through protein degradation. Compared to other chemicals, FDA-approved drugs have been well characterized in toxicity and efficacy, making them more appreciated for drug repurposing; thus, we determined to screen for the chemicals from this library. To perform the drug screening, we first established a cell-based screening platform to measure the abundance of PreS2-LHBS. Specifically, we adopted a protein stability reporter system in 293T cells [20]. This system evaluated the PreS2-LHBS abundance by two fluorescent signals: EGFP, tagged with PreS2-LHBS; DsRed, an internal control, synthesized from the same mRNA of EGFP (Figure 1A). The established reporter stable cell line expressing PreS2-LHBS in 93.4% of the cell population was used in this study (Figure S1). Next, we individually challenged cells with one chemical from the FDA-approved drug library (1068 drugs) and assessed fluorescent signals and cell viability. According to Chebyshev’s inequality, around 90% of data will locate within 3-fold of standard deviation (SD) around the mean without any model assumption. Therefore, we considered the drug significantly reduced PreS2-LHBS when the fluorescent signal was beyond 3 SD from the mean (Figure 1A). We selected drugs that significantly reduced fluorescent signal without affecting cell viability as our initial candidates. From a collection of 1068 drugs, 11 drugs hit this criterion (Figure 1B). These drugs reduced around 40% to 60% of fluorescence signal but maintained over 80% cell viability (Figure 1C). In addition, hierarchical clustering analysis revealed that screen hits were in the same cluster, indicating that we selected a group of drugs with a similar mechanism of action on PreS2-LHBS, strengthening our selection criterion. (Figure 1D). We treated SNAP-tagged PreS2-LHBS stable hepatocyte cell line (NeHep-S2) with selected drugs and evaluated the expression level of PreS2-LHBS by immunoblotting. We confirmed that ABT199, Rapamycin, Bupivacaine, and Deferoxamine reduced intracellular PreS2-LHBS (Figure 2A).

**Figure 1.**
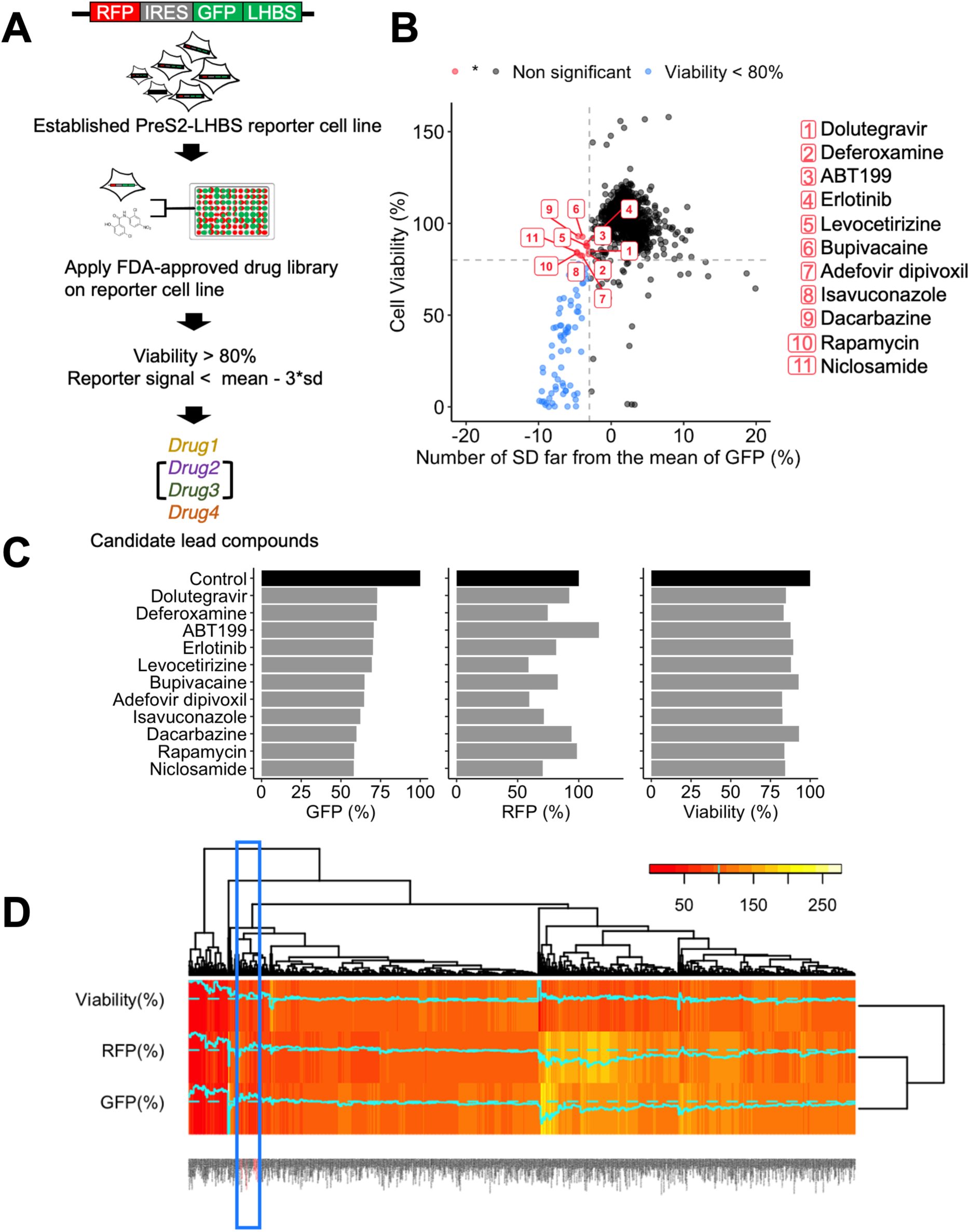
Repurposing FDA-approved drug to target PreS2-LHBS through protein degradation. (A) Schematic overview of FDA-approved drug library screening. (B) Scatter plot of screening result. PreS2-LHBS reporter cells were treated with 10uM of one drug. Each point represents the value of cells viability and the number of standard deviations (SD) far from the mean in each treatment. The vertical line indicates 3 SD below the mean, and the heretical line indicates 80% cell viability. The blue points indicated the drug below 80% viability, and the red points indicates screen hits. (C) The intensity of three measurements relative to solvent control (DMSO) in FDA-approved drug screening. (D) Hierarchical clustering analysis of screening results. Each column is a drug clustered by statistical distance according to its intensity in three measurements. The blue box highlights the cluster containing the selected drugs (colored by red). Color key shows the range of intensity in three measurements. Cyan line indicates the intensity value across each drug.

**Figure 2.**
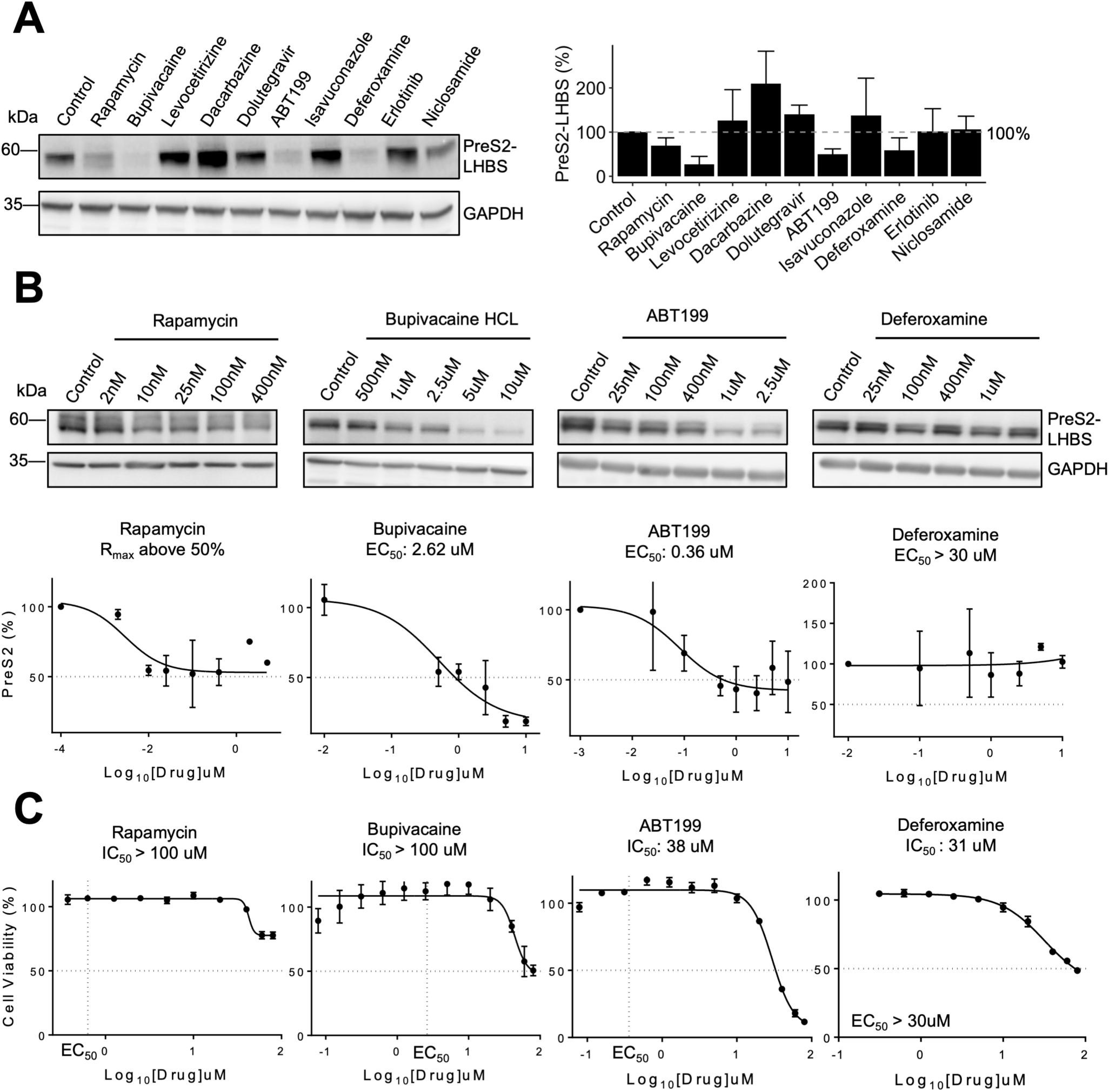
Efficacy of the repurposed drug. (A) Verification of selected drug. NeHep-S2 cells were subjected to 10uM selected drug, and the protein expression was analyzed by immunoblotting. Left panel: representative blot. Right panel: bar chart shows the expression normalized to solvent control (DMSO) and loading control (GAPDH). (B) EC_50_ of the repurposed drug. NeHep-S2 cells were treated with the indicated concentration of the repurposed drug, and the protein expression was analyzed by immunoblotting. Upper panel: representative blot. Bottom panel: The absolute EC_50_ was calculated by a three-parameter logistic function fitted to the concentration against the expression. The expression was normalized to solvent control (DMSO) and loading control (GAPDH). (C) The cytotoxicity of the repurposed drug was determined by cell viability. Six thousand NeHep-S2 cells were seeded in 96 plates and subjected to repurposed drug with indicated concentration for 24hr to measure the cell viability. The relative IC50 was calculated by a three-parameter logistic function fitted to the concentration against the viability. The viability was normalized to solvent control (DMSO) treated cells. The vertical line shows the EC50 of each drug, and the heretical line shows 50% cell viability. Error bar show SEM.

### ABT199 reduces intracellular expression of PreS2-LHBS

To assess the efficacy of repurposed drugs on reducing PreS2-LHBS, we evaluated the 50% effective concentration (EC_50_) and the maximal reduction percentage (R_max_). To this end, NeHep-S2 cells were treated with each drug in serial dilution. We found that ABT199, Bupivacaine, and Rapamycin, but not Deferoxamine, reduced intracellular PreS2-LHBS in a dose-dependent manner. Of three effective drugs, ABT199 showed the lowest EC_50_; ABT199 and Bupivacaine showed R_max_ higher than 80% (Figure 2B). Notably, we observed that the reduction of PreS2-LHBS (>30%) occurred upon treatment with all three drugs in nanomolar concentration. Furthermore, these drugs showed no cytotoxicity (∼100% cell viability) in their effective concentration (Figure 2C), indicating that the efficacy of reducing PreS2-LHBS was not due to cell death. Because ABT199 was the most effective drug and can be taken orally, we focused on ABT199 in the following study.

To better understand how ABT199 reduced PreS2-LHBS, we examined the kinetics of PreS2-LHBS reduction upon treatment of ABT199 over time. NeHep-S2 cells were treated with ABT199 and were followed the expression by immunoblotting. We found that ABT199 significantly reduced PreS2-LHBS within 6hr (>50%), whereas the internal control, BCL2, was not affected (Figure 3A). Next, we tested if ABT199 affects the PreS2-LHBS protein half-life, a hallmark for estimating intracellular protein stability. To measure the protein half-life, we blocked the protein synthesis by cycloheximide in the presence or absence of ABT199. Strikingly, fitting the expression with non-linear regression showed that ABT199 reduced the PreS2-LHBS protein half-life by 54%, shirking from 5.7hr to 2.6hr (Figure 3B), suggesting PreS2-LHBS tends to be degraded in the presence of ABT199. Finally, we verified the efficacy of ABT199 in HuH7 and HepG2 infected with adenovirus carrying a complete wild-type HBV genome (AdHBV) [21]. As expected, ABT199 reduced the intracellular LHBS in HuH7 and HepG2 by around 40% (Figure 3C). Accordingly, our data indicate that ABT199 effectively reduces either wild-type or PreS2-LHBS in the context of the complete HBV genome.

**Figure 3.**
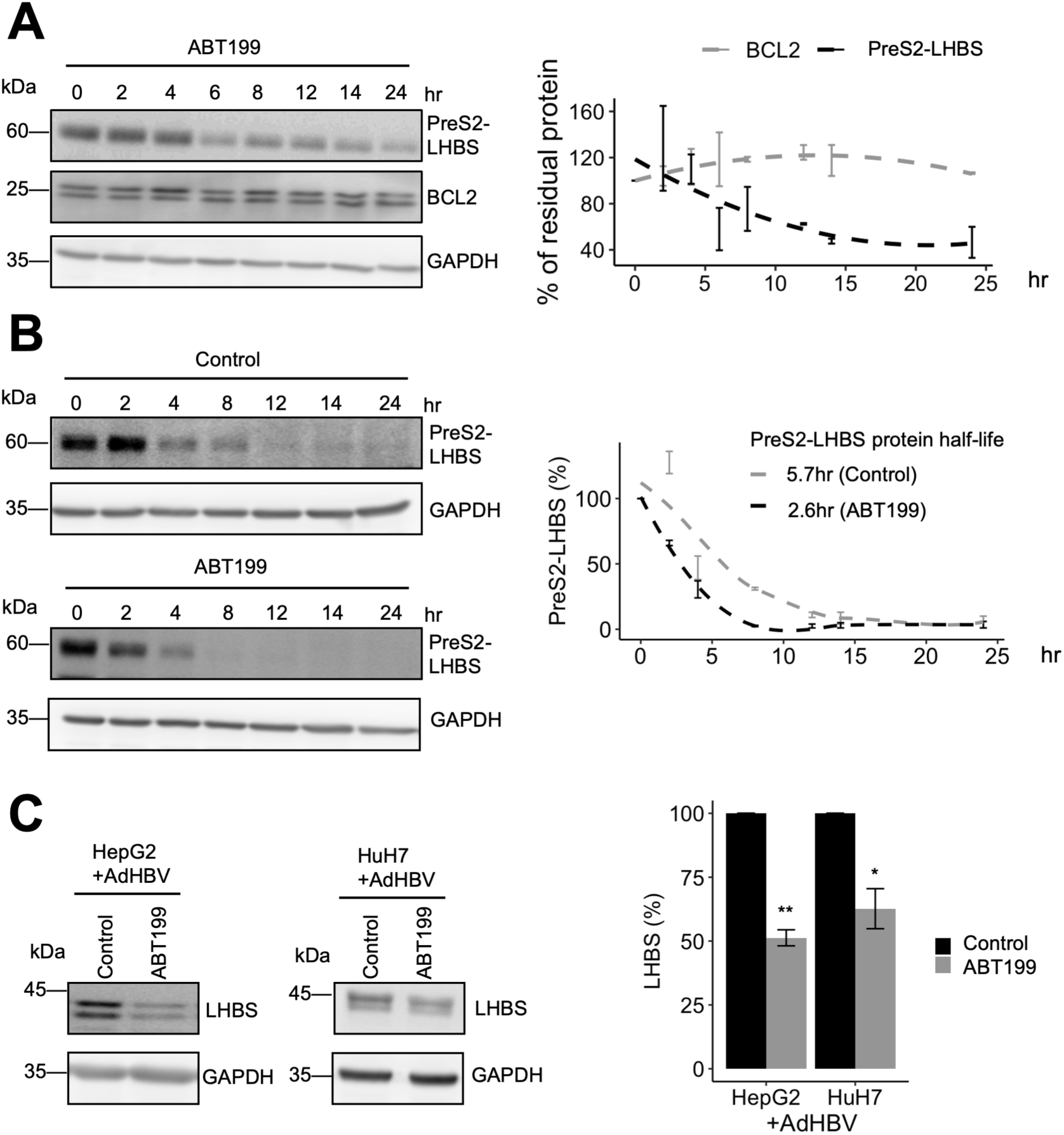
ABT199 significantly reduces PreS2-LHBS. (A) The reduction kinetics of PreS2-LHBS induced by ABT199. NeHep-S2 cells were treated with ABT199 (1uM) and were chased for protein expression at indicated time points by immunoblotting. BCL2 served as an internal control. Left panel: representative blot. Right panel: The graph shows the nonlinear regression fitted to the time point against the protein expression. The protein expression was normalized to the initial time point (0hr) and loading control (GAPDH). (B) Measure of PreS2-LHBS protein half-life. NeHep-S2 cells were treated with cycloheximide (10 ug/ml) in the presence or absence of ABT199 (1uM). The protein expression was chased for indicated time points by immunoblotting. Left panel: representative blot. Right panel: The protein half-life was calculated by nonlinear regression fitted to the time point against the expression. The expression was normalized to the initial time point (0hr) and loading control (GAPDH). (C) The reduction of LHBS induced by ABT199 was verified using AdHBV infected cells. After 48hr post-infection with adHBV at 100 MOI, HuH7 and HepG2 cells were treated with ABT199 (1uM) or solvent control (DMSO) for 24hr. Cells were then analyzed by immunoblotting. Left panel: representative blot. Right panel: bar chart shows the expression normalized to solvent control (DMSO) treated cells and loading control (GAPDH). Error bar showed SEM. **P < 0.005, *P < 0.05, determined by two-tailed, unpaired Student’s t-test.

### ABT199 induces PreS2-LHBS degradation through microautophagy

From the previous section, we hypothesized that ABT199 reduced PreS2-LHBS by activating cellular pathways; thus, we aimed to identify the cellular pathways that control the protein level of PreS2-LHBS. To study whether PreS2-LHBS transcription is affected by ABT199, NeHep-S2 cells were treated with ABT199 or solvent control (DMSO), and the mRNA level was analyzed by quantitative real-time PCR. We found no significant difference in PreS2-LHBS mRNA level (Figure 4A). Thus, ABT199 does not affect the transcription of PreS2-LHBS. Next, we tested whether ABT199 directly interacts with LHBS to trigger the degradation. To this end, we incubated purified LHBS protein with ABT199 or solvent control (DMSO) and measured the interaction by microscale thermophoresis (MST). Compared to solvent control, the interaction between LHBS and ABT199 showed no correlation with ABT199 concentration (Figure S2), suggesting ABT199 does not directly interact with LHBS. Thus, we sought to explore the degradation pathway activated by ABT199.

**Figure 4.**
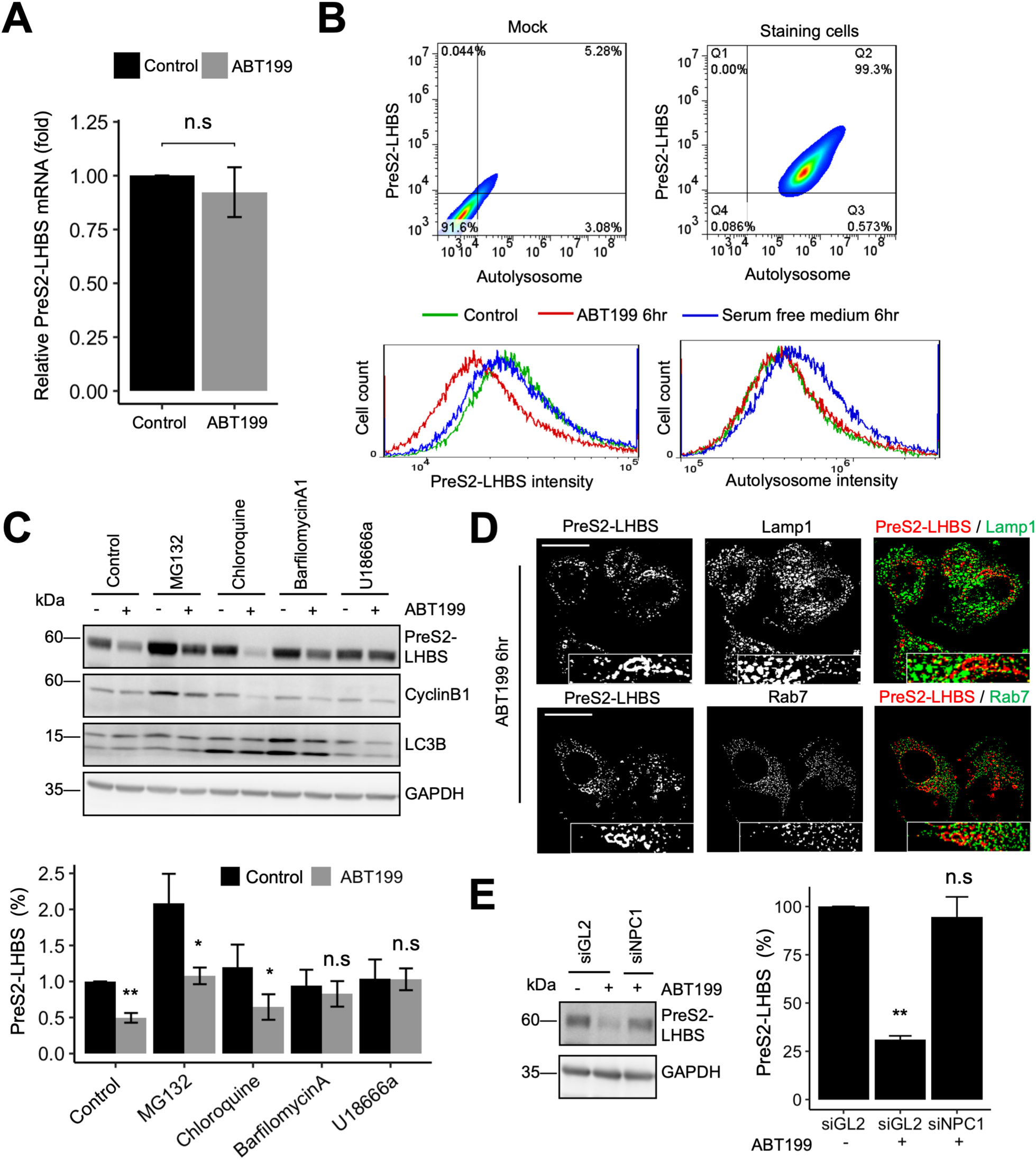
ABT199 induces PreS2-LHBS degradation through microautophagy. (A) NeHep-S2 cells were treated with ABT199 (1uM) or solvent control (DMSO), and the mRNA level was measured by qPCR. The bar plot shows the mRNA level relative to ACTB mRNA and normalized to solvent control (DMSO) treated cells. (B) Examined the activation of macroautophagy. Upper panel: the intensity profile of NeHep-S2 cells staining for PreS2 mutant LHBS and autolysosome. Bottom panel: after incubated with solvent control (DMSO), ABT199 (1uM), or in serum-free medium for 6hr, ten thousand stained cells were analyzed by flow cytometry. The density plot shows the cell counts for the intensity of PreS2-LHBS and autolysosome in each condition. (C) Investigation of ABT199-induced pathway for PreS2 mutant LHBS degradation. Nehep-s2 cells were treated with indicated inhibitors with or without ABT199 (1uM), and the protein expression was analyzed by immunoblotting. The level of CyclinB1 and LC3B served as a positive control for inhibitor treatment. Upper panel: representative blot. Bottom panel: bar plot shows the expression normalized to solvent control (DMSO) treated cells and loading control (GAPDH). (D) NeHep-S2 cells were treated with ABT199 (1uM) for 6hr, and the colocalization of PreS2 mutant LHBS, Lamp1, and Rab7 were investigated by immunofluorescent staining. Scale bar donates 20um. (E) Depletion of NPC1 rescued ABT199-induced PreS2 mutant LHBS degradation. NeHep-S2 were knockdown by transfection with a control siRNA or NPC1-specific siRNA, and the protein expression was examined after 48hr post-transfection by immunoblotting. Left panel: representative blot. Right panel: bar plot shows the expression normalized to loading control (GAPDH) and cells transfected with control siRNA. Error bar shows SEM. **P < 0.005, *P < 0.05, n.s, non-significant determined by two-tailed, unpaired Student’s t-test.

ABT199 is a BH3 domain analog that selectively attaches to BCL2 and inhibits its activity. Previous studies have shown that dissociation of Beclin-1 and BCL2 can initiate macroautophagy, a well-studied protein degradation pathway [22]. Therefore, we tested whether ABT199 may activate macroautophagy. We examined the macroautophagy signal, the localization between PreS2-LHBS and autolysosome, and the phenomena after depleting BCL2 and/or Beclin1 in ABT199 treated cells (Figure 4A, S3, S4); however, we found no evidence supporting the activation of macroautophagy. Thus, we turned to investigate all possible degradation pathways. The protein degradation pathway can be classified into: proteasomal degradation and lysosomal degradation, where lysosomal degradation can be further classified into: macroautophagy, microautophagy, and chaperone-mediated autophagy. To identify key degradation pathways, we treated cells with each pathway inhibitor in the presence or absence of ABT199 and examined the expression of PreS2-LHBS. We found that inhibition of either lysosomal degradation or microautophagy abolished ABT199-induced degradation, indicating that microautophagy is involved in regulating the intracellular stability of PreS2-LHBS (Figure 4C). Notably, ABT199-induced degradation was also abolished upon depletion of NPC1, which inhibited microautophagy’s dynamic (Figure 4E). Immunofluorescent staining showed that PreS2-LHBS was recruited to a ring-shaped structure, and the distribution was in close proximity of the lysosomal marker Lamp1 and MVB marker Rab7 upon treatment of ABT199 (Figure 4D). In addition, we performed a purification screening following mass spectrometry analysis and identified HSC70, an essential chaperone mediating microautophagy [23], significantly interacted with PreS2-LHBS (Figure S5). Taken together, our data suggest that ABT199 activated microautophagy to degrade PreS2-LHBS.

### Long-term inhibition of PreS2-LHBS by ABT199 reverses oncogenic effect in PreS2-LHBS expressing hepatocytes

To understand the long-term impact of reducing PreS2-LHBS, we tested if LHBS-induced oncogenic events such as increased DNA damage, mitotic defect, chromosome hyperploidy, and oncogenesis [9] can be reversed by ABT199. To evaluate the DNA damage, we stained ABT199-treated and untreated NeHep-S2 cells for phosphorylated H2AX (pH2AX), a DNA damage marker, and calculated the percentage of pH2AX positive cells in the population. Strikingly, compared to control, we found that ABT199 significantly reduced DNA damage in PreS2-LHBS cells by 2.1-fold (Figure 5A). In addition, long-term treatment (3weeks) of ABT199 not only significantly reduced DNA damage but showed a lower variation of positive percentage, suggesting that persistent suppression of PreS2-LHBS can restore DNA damage much effectively. Next, to monitor mitotic failure, we followed the mitotic progression of NeHep-S2 cells by time-lapse imagining and classified mitotic events according to their division results (Figure 5B, left panel). Compared to control cells, we found that ABT199 significantly reduced 45% of mitotic failure in NeHep-S2 cells (Figure 5B, right panel). Furthermore, analysis of mitotic failure revealed that the reduction came from the decrease of cytokinesis failure (Figure 5B, right panel), which is identical to the observation while depleting LHBS in our previous study [9].

**Figure 5.**
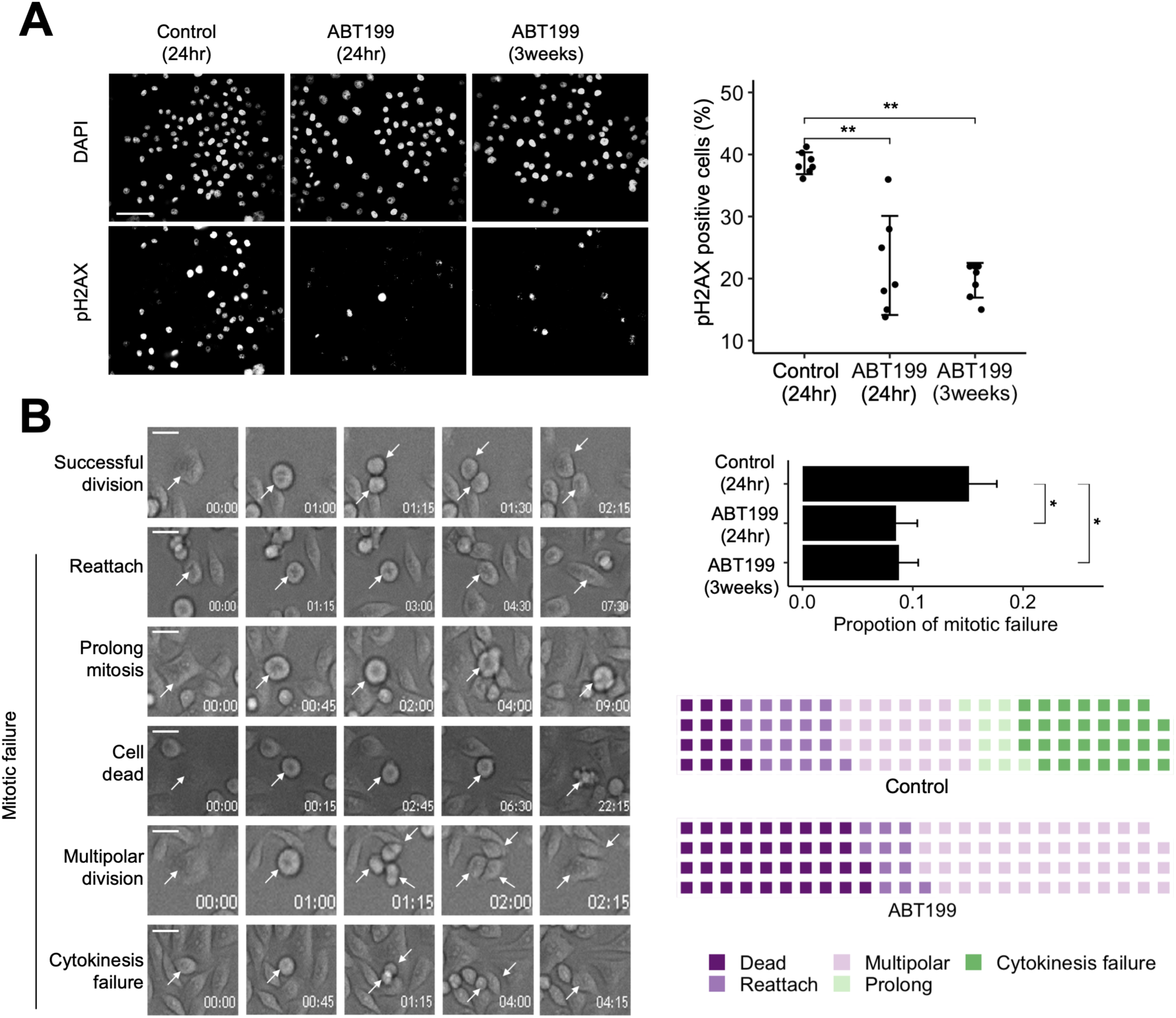
Inhibition of PreS2-LHBS by ABT199 reduces oncogenic events in PreS2-LHBS expressing hepatocytes. (A) NeHep-S2 cells were treated with solvent control (DMSO) or ABT199 (1uM) for the indicated time, and the DNA damage was determined by staining for pH2AX. Left panel: representative image. Right panel: graph shows the percentage of positive cells. Scale bar donates 100um. (B) NeHep-S2 cells were treated with solvent control (DMSO) or ABT199 (1uM) for the indicated time, and the mitotic events were followed by time-lapse imaging for 24hr. Left panel: classification of mitotic event. Upper right panel: bar chart shows the proportion of mitotic failure in the division events. Bottom right panel: waffle plot resolves the composition of mitotic failure in each treatment. Each square represents one percentage in failure events. Scale bar donates 10um. (C) NeHep-S2 cells were treated with ABT199 (1uM) or solvent control (DMSO) for 3weeks, and the chromosome numbers in the cells were calculated by chromosome spreading. Left panel: representative image. Middle panel: the distribution of chromosome numbers in the cells is showed by histogram. Right panel: scatter plot displays the chromosome numbers in each calculated cell from three experiments. Scale bar donates 20um. Error bar showed SEM. **P < 0.005, *P < 0.05, n.s., non-significant determined by two-tailed, unpaired Student’s t-test.

We next examined if ABT199 can reduce LHBS-induced hepatocyte hyperploidy. We treated cells with ABT199 for 3weeks and calculated the chromosome numbers by chromosome spread. NeHep-S2 cells displayed chromosome numbers of 57 on average, showing a right-skewed distribution. In comparison, long-term treatment of ABT199 significantly reduced chromosome number to an average of 50 that reverted to a normal distribution (Figure 6A). Specifically, the number of cells containing more than 55 chromosomes was significantly reduced by ABT199, suggesting the persistent reduction of PreS2-LHBS reduced hepatocyte hyperploidy cells. Finally, we test if long-term treatment of ABT199 may inhibit LHBS-induced oncogenesis. Soft agar colony formation assay is a gold standard method to evaluate the oncogenic property of transformed cells; only transformed cells show anchorage-independent growth on 3-dimensional soft agar [24]. We seeded HepAD38 cells (a hepatoma cell line expressing all viral proteins including wild-type LHBS [25]) in the 3-dimensinal soft agar plates and treated cells with solvent control (DMSO), ABT199, entecavir (ETV, a conventional antiviral drug targeting HBV reverse transcription), or co-treated with ABT199 and ETV, respectively. Interestingly, only treatment of ABT199 significantly inhibited colony formation (Figure 6B). Compared to ABT199, ETV did not significantly inhibit colony formation; however, the colony number and size were further reduced by co-treatment with ABT199, suggesting the oncogenesis can be constrained by targeting LHBS. In summary, these data show that persistently reducing PreS2-LHBS by ABT199 could consequently revere the oncogenic effects caused by PreS2-LHBS.

**Figure 6.**
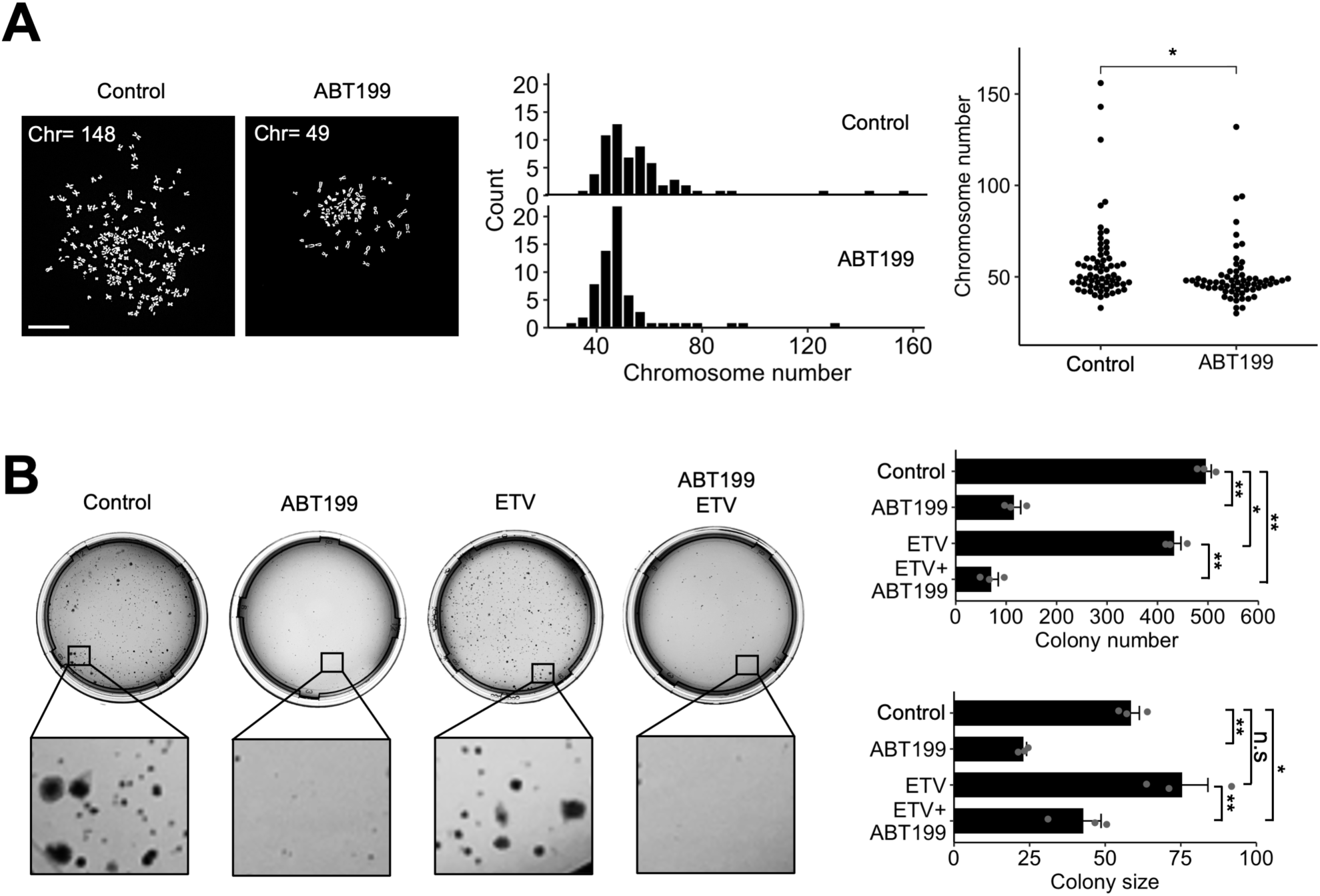
Long-term inhibition of LHBS constrain HBV-mediate oncogenesis. (A) NeHep-S2 cells were treated with ABT199 (1uM) or solvent control (DMSO) for 3 weeks, and the chromosome numbers in the cells were calculated by chromosome spreading. Left panel: Number of counted chromosome number in the representative image is showed. Middle panel: the distribution of chromosome numbers in the cells is showed by histogram. Right panel: scatter plot displays the chromosome numbers in each calculated cell from three experiments. Scale bar donates 20um. (B) HepAD38 cells were cultured in soft agar plates containing solvent control (DMSO), ABT199 (2uM), or ETV (50nM) as indicated for 3 weeks. The total colony number and size were calculated by ImageJ. Left panel: representative image. Right panel: bar plot shows the colony number and size in each treatment. Error bar showed SEM. **P < 0.005, *P < 0.05, n.s., non-significant determined by two-tailed, unpaired Student’s t-test.

## Discussion

In this study, we repurposed several new drug candidates to target intracellular PreS2-LHBS. According to HBV drug watch, most of the developing anti-HBV drugs take action on HBV viral polymerase and capsid, whereas only two chemicals target extracellular HBsAg in clinical development [26]. To our best knowledge, inhibitors that specifically target intracellular surface antigens are not available. As such, these candidates represent the first chemical inhibitors of intracellular PreS2-LHBS.

Targeting viral surface antigen faces several technical difficulties. First, surface antigen is an envelope structural protein with no enzyme activity. Therefore, it is not possible to develop an *in vitro* functional enzyme assay for the selection of chemical inhibitors. Second, surface antigen can be translated from either HBV cccDNA or integrated genome, and the integration of HBV DNA occurs within three days post-infection [27], making it challenging to eliminate surface antigen at the DNA level. In addition, RNA interference is likely inefficient to eliminate surface antigen in the context of a whole liver. A recent study showed that siRNA, designed using a complete HBV sequence, could not efficiently reduce surface antigens in HBeAg-negative patients due to the increased integrated DNA and viral sequence mutation following CHB progression [17]. As an alternative approach, we tested whether any inhibitor can effectively target the PreS2-LHBS at the protein level. Activating the cellular degradation pathway to promote the clearance of toxic protein has been described in neurodegenerative diseases, such as Alzheimer’s disease [28]. These neurodegenerative diseases share a common feature: accumulation of non-function protein, similar to CHB that accumulates PreS2-LHBS in hepatocytes. With this central ideal, we adopted the protein stability platform and identified Rapamycin, Bupivacaine, and ABT199 that induced selected degradation of intracellular PreS2-LHBS. Interestingly, all these screen hits are linked to different processes of autophagy. Rapamycin is a well-studied mTOR inhibitor that triggers autophagy and ubiquitin-proteasome system [29]; Bupivacaine has been reported to induce protein degradation in muscle cells [30,31] and also induce autophagy in lung cancer cells [32]; ABT199 is a BCL2 inhibitor that has the potency to regulate apoptosis and autophagy pathway [33]. These data imply that manipulating protein-specific degradation is a plausible approach to inhibit PreS2-LHBS.

In this study, we found that microautophagy is activated by ABT199. Endosomal microautophagy occurs during the formation of MVB to deliver the protein into the late endosome/MVB vesicle and degrade the content either directly in the late endosome or by the fusion with the lysosome. This process requires the ESCRT component and HSC70, an essential chaperone, to sequester protein into the endosome membrane [23]. Notably, we also identified that HSC70 interacted with PreS2-LHBS from purification screening upon ABT199 treatment (Figure S6). How exactly ABT199 activates microautophagy is not clear. By searching the SwissTargetPrediction [34], mTOR is listed as a potential interaction partner of ABT199 with a high prediction probability. mTOR has been reported to regulate microautophagy [35,36], suggesting a possible role of microautophagy in the stability control of PreS2-LHBS. We suggest the interaction with HSC70, and the implication of endosomal transport may contribute synergistically to the degradation of PreS2-LHBS in the presence of ABT199.

Whether the side effect of ABT199 is acceptable for CHB treatment? ABT199 is a BH3 domain analog, mainly used to treat chronic lymphocytic leukemia by inhibiting BCL2 function [37]. Tumor lysis syndrome, caused by a large number of tumor cell death, is the major precautions side effect in treated patients [38]. In this study, the EC_50_ of ABT199 in inhibiting PreS2-LHBS was much lower than the IC_50_ of cell viability, suggesting that ABT199 may inhibit PreS2-LHBS without affecting general cell viability. Previous studies have shown that the inhibition of BCL2 can disrupt HBV-induced calcium imbalance, drug resistance, and viral replication [39,40]. We also did not detect cytotoxicity while depleting BCL2 in PreS2-LHBS expressing cells (Figure S3). These data imply that (1) depletion of BCL2 is not cytotoxic to HBV-positive hepatocytes and (2) inhibition of BCL2 may block HBV through multiple intracellular pathways, including the induction of microautophagy as shown in this study, and the reversion of calcium imbalance, drug resistance, and viral replication.

Finally, we showed that long-term inhibition of PreS2-LHBS by ABT199 can significantly decrease DNA damage, cytokinesis failure, hyperploidy, and oncogenesis. Surface antigen has been proposed to induce ER stress, chromosome instability, and HCC [15,41,42]. Our results highlight the possibility of reversing these oncogenic effects by reducing surface antigen. Interestingly, hyperploidy population seems to be an inflexible consequence but also can be reduced by long-term inhibiting PreS2-LHBS. A study has shown that hepatocytes can undergo ploidy reversal to divide into different genome copies containing hepatocytes [43]. Our result suggests that PreS2-LHBS is required for maintaining the hyperploidy population. Due to the time required for ploidy reversal [43,44], we notice that long-term inhibition of Pres2-LHBS is the key to restricting oncogenesis. Current methods such as CRISPR/Cas9 gene editing may be applied to long-term inhibition of PreS2-LHBS. Studies have shown that both integrated and cccDNA can be targeted by CRISPR/Cas9 system, resulting in the inhibition of viral product [45–47]. However, CRISPR/Cas9 system requires the pre-modified cells expressing Cas9 protein, which might be challenging for clinical application. In addition, eliminating all the HBV genomes in the infected hepatocytes is also tricky. Although the long-term efficacy of ABT199 needs further investigation in different advanced models, and there might be other methods to achieve long-term inhibition of PreS2-LHBS, ABT199 is an on-broad method for targeting PreS2-LHBS. Importantly, our result proves the concept that long-term reduction of PreS2-LHBS is a plausible therapeutic option to prevent HBV-induced oncogenesis. Whether such an approach may advance the complete elimination of HBV in patients is still awaiting further examinations.

## Materials and Methods

### Molecular cloning and cell culture

To produce PreS2-LHBS reporter construct, PreS2-LHBS sequence (deletion nucleotides 2-55) was cloned into the protein stability construct after the c-terminal of EGFP by In-Fusion cloning (Takara). The protein stability construct was a kind gift from Dr. Hsueh-Chi Sherry Yen, Academia Sinica, Taiwan [20]. To generate protein stability reporter cell line, 293T cells were transfected with the protein stability construct carrying PreS2-LHBS. Transfected cells were selected by 2 µg/ml puromycin for two weeks and further selected by cell sorting according to their protein stability signal. Selected cells were limit diluted into 96 well plates to obtain the single clone. Protein stability signal and PreS2-LHBS expression of single clones were then confirmed by immunoblotting and flow cytometry. The immortalized hepatocyte cell line, NeHepLxHT, was obtained from the American Type Culture Collection. NeHepLxHT stably expressing SNAP-tagged PreS2-LHBS (NeHep-S2) were established as described previously [15]. NeHepLxHT was maintained in Dulbecco’s Modified Eagle’s Medium/Hams F-12 50/50 Mix (Corning) supplied with 15% FBS (Biological industries), 1% penicillin-streptomycin (Corning), 0.1% ITS universal cell culture supplement premix (BD), and 20 ng/ml epidermal growth factor (BD). 293T, HuH7, and HepG2 were maintained in Dulbecco’s modified Eagle’s medium (DMEM, Corning) supplied with 10% FBS and 1% penicillin/streptomycin. All lines were maintained in a humidified atmosphere supplied with 5% CO2 at 37 °C.

### Reagents

The following reagents were used: DMSO (as solvent control, sigma), ABT199 (1uM, Selleckchem), cycloheximide (10mg/ml, MDBio), MG132 (10uM, CAYMAN chemical), Chloroquine (10uM, sigma), BarfilomycinA1 (1uM, Selleckchem), and U18666a (10ug/ml, Biovision).

### FDA-approved drug library screening and cell viability assay

The FDA-approved drug library was purchased from TargetMol (No. L1010). PreS2-LHBS reporter stable cells were seeded in 96 well plated and treated with 10uM of one drug for 24hr. The culture medium was removed, and the cells were washed with PBS two times. The EGFP and DsRed signal were measured by a microplate reader (BioTek Synergy HTX) with the excitation wavelength: 485/20, 540/35; emission wavelength: 528/20, 600/40, respectively. CellTiter-Glo (Promega) was used to measure the viability according to the manufacturer’s instructions. The luminescence signal was measured by a microplate reader (BioTek Synergy HTX). All readout was normalized to solvent control (DMSO)

### Immunoblotting

Cells were lysed in RIPA buffer and were centrifuged at 4 °C, 13000 rpm for 15 minutes to collect the total protein. Total protein was denatured in laemmli buffer at 100°C for 15 minutes and was transferred to the PVDF membrane after being separated by SDS-PAGE. The membranes were incubated with primary and secondary HRP-conjugated antibodies in PBST (5% milk and 0.1% Tween-20) to detect the protein of interest by Western Lightning Plus ECL (PerkinElmer) and ImageQuant LAS 4000 imaging system (GE). Protein expression was quantified by ImageQuant TL software and normalized to treatment control and loading control. The following antibody were used: mouse anti-LHBS/PreS1 (7H11, a kind gift from Professor Ning-Shao Xia, Xiamen University, China 18), mouse anti-BCL2 (sc7382, Santa Cruz), rabbit anti-LC3B (GTX127375, GeneTex), rabbit anti-cyclinB1 (GTX100911, GeneTex), rabbit anti-GAPDH (GTX100118, GeneTex), goat anti-rabbit IgG HRP-conjugated (H+L, 111-035-003, Jackson ImmunoResearch), and goat anti-mouse IgG HRP-conjugated (H+L, 115-035-003, Jackson ImmunoResearch).

### RT-qPCR

The total mRNA was extracted from the cells by REzol C&T (PT-KP200CT) using a standard RNA extraction protocol. The mRNA was reverse transcribed into cDNA by SuperScript™ IV VILO™ Master Mix (11756050, ThermoFisher) with an oligo(dT) primer. The quantitative PCR was performed with SensiFAST SYBR Lo-ROX Kit (Bioline), 0.2uM forward and reveres primer, 1ul of 1:10 diluted cDNA in 10ul reactions, running on AriaMx Real-Time PCR system (Agilent Technologies) using a standard thermal cycle program with 25 s at 62 °C for annealing and 30 cycles. Primer used were against PreS1/LHBS (forward: 5’-CCCGATCATCAGTTGGACCC-3’, reverse: 5’-TGGTCAATATGCCCTGAGCC-3’), ACTB (forward: 5’-CATTGCTGACAGGAT-GCAGAAGG-3’, reverse: 5’-TGCTGG AAGGTGGACAGTGAGG-3’).

### Immunofluorescence staining and epifluorescence imaging

Cells were cultured on glass coverslips and treated with the indicated chemical for indicated time point before fixation. Cells were then fixed with 4% formaldehyde, 1 μM MgCl2, 20 uM PIPES, 10 μM EGTA, and 0.2% Triton X-100 and washed with PBST (0.1% Tween-20). Coverslips were incubated with primary antibody for 1hr and secondary antibody for 30 mins in PBST (5% FBS and 0.1% Tween-20). Antibodies used were rabbit anti-Lamp1 (9091S, Cell signalling), rabbit anti-Rab7 (ab137029, Abcam), mouse anti-PreS2/LHBS (7h11), rabbit anti-pH2AX-Ser139 (JBW301, Merck), donkey anti-rabbit IgG Alexa Fluor 488-labeled (H+L, A21206, Invitrogen), and donkey anti-rabbit IgG Alexa Fluor 594-labeled (H+L, A21207, Invitrogen). DNA was stained by DAPI. Coverslips were mounted with Vectashield antifade mounting medium (Vector Laboratories), and the images were acquired by epifluorescence microscopy (Leica DMI6000). For analysis of macroautophagy, cells were cultured on glass coverslips and treated with either 10uM chloroquine (negative control) or 1uM ABT199 for the indicated time, following staining for PreS2-LHBS and autolysosome by SNAP-Cell® TMR-Star (S9105S, NEB) and DALGreen (D675-10, Dojindo), respectively. Cells were then fixed with 4% formaldehyde and mounted with Vectashield antifade mounting medium (Vector Laboratories). Images were acquired by epifluorescence microscopy (Leica DMI6000).

### Live cell microscopy

Cells were grown in chamber slides (Ibidi) at 80% confluent. Time-lapse images were acquired at 15 min intervals for 24 h by epifluorescence microscopy (Leica DMI6000) equipped with Andor Luca R EMCCD camera (Andor Technology) and a motorized stage supplied with 5% CO2 at 37°C. Mitotic cells were followed once the cells rounded up and were classified by their destination. At least 100 cells were counted in each condition from three independent experiments. Images were analyzed by MetaMorph software (Molecular Devices).

### Chromosome spreads

Cells were synchronized to M-phase by treating with 10ng/ml nocodazole overnight and were then collected by mitotic shake-off. Collected cells were incubated in 75 mM KCl at room temperature for 20 min and were replaced with Carnoy’s solution (freshly prepared, 3:1 of ice-cold methanol and glacial acetic acid) for 30 min on ice. Cells were dropped on glass slides to spread the chromosome. Chromosomes were stained with DAPI and were imaged by epifluorescence microscopy (Leica DMI6000) to count the total chromosome number in each cell.

### Colony formation assay

Soft agar plates were generated by a bottom layer and an upper layer. Bottom layer contained DMEM-F12, 1.2% agar. Upper layer contained 600 of HepAD38 cells, DMEM-F12, and 0.3% agar. Solvent control (DMSO), ABT199 (2uM), ETV (50nM) were added into both layers as indicated. Cultured medium containing corresponded chemical were add into the dish each weeks. After culturing for 3 weeks, cells were stained with 10% trypan blue. The colony number and size were then analyzed by ImageJ.

### AdHBV infection

Adenoviral HBV gene transfer (AdHBV), carrying a 1.3-fold HBV genome, was a kind gift from Dr. Li-Rung Huang, National Health Research Institutes, Taiwan [21]. To examine the efficacy of ABT199, cells were seeded in 6cm at 60% confluent and were infected with 200MOI AdHBV for two days. The infected cells were then treated with 1uM ABT199 for 24hr and assayed the protein expression by immunoblotting.

### Flow cytometry analysis

For analysis of macroautophagy activation, cells were double-stained for autolysosome and PreS2-LHBS by DALGreen and SNAP-Cell® TMR-Star after being treated with the indicated condition. Cells were resolved by flow cytometer (CytoFLEX, BECKMAN COULTER). At least one thousand cells were acquired in each experiment. FCS3.0 files were converted by FCS extract (1.02) and were imported into FlowJo (v.10.6.2) for further analysis and figure generation. FSA-A and SSC-A channels were used to gating the analyzed cells. FITC-A and PE-A channels were used to analyze the level of autolysosome and PreS2-LHBS, respectively.

### siRNA transfection

RNA interference (RNAi) was performed by siRNA transfection using DharmaFECT 1 Transfection Reagent (Dharmacon) according to the manufacturer’s instruction. siRNA (on-target, smart pool) was purchased from Dharmacon. The following siRNA used were against BLC2 (5′-GGGAGAACAGGGUACGAUA-3′, 5′-GAAGUACAUCCAUUAUAAG-3′, 5′-GGAGGAUU-GUGGCCUUCUU-3′, 5′-UCGCCGAGAUGUCCAGCCA-3′), BECN1 (5′-GAUACCGA-CUUGUUCCUUA-3′, 5′-GGAACUCACAGCUCCAUUA-3′, 5′-CUAAGGAGCUGCCGUU-AUA-3′, 5′-GAGAGGAGCCAUUUAUUGA-3′) and control (5′-CGUACGCGGAAUACUUCGAdTdT-3′)

### Microscale thermophoresis analysis

The purified HBsAg was purchased from ProSpec (hbs-872-c) and was labeled by RED-NHS dye (monolith protein labeling kit, MO-L011, NanoTemper) according to the manufacturer’s instruction. The labeled protein was 1:1 incubated with a series dilution of ABT199 (4 uM to 0.122 nM) to perform the microscale thermophoresis experiments in Monolith NT.115 (NanoTemper) with the setting of 40% MST power and 40% excitation power at 25 °C. The data were analyzed by the default software.

### Immunoprecipitation followed by quantitative mass-spec analysis

To pull down PreS2-LHBS, cells were lysate in Pierce buffer (25 mM Tris-HCl pH 7.4, 150 mM NaCl, 1 mM EDTA, 1% NP-40 and 5% glycerol) containing either solvent control (DMSO), 1uM ABT199, or 1uM ABT199 and u18666a. Cell lysates were centrifuged at 4 °C, 13000 rpm for 15 minutes to collect the total protein. The total lysate was incubated with the protein G dynabeads (10004D, Invitrogen) conjugated with mouse anti-PreS1 antibody (7h11) or mouse IgG (control, sc-2025, Santa Cruz) overnight, and the PreS2-LHBS interacted complex was pulled down according to the manufacturer’s instruction. The immunoprecipitation products were eluted from the beads by TFA. The quantitative mass-spec analysis was performed by Biotools, Superlab using an LC apparatus coupled with 2D linear ion trap mass spectrometer (Orbitrap Elite, Thermo Fisher Scientific) after labeling in-solution digested (sequencing-grade modified porcine trypsin, Promega) immunoprecipitation product with iTRAQ reagent. The protein identification and quantification were performed by Proteome Discoverer software (version 1.4, Thermo Fisher Scientific)

### Quantification and statistical analysis

For analyzing FDA-approved drug library screening, the fluorescence intensity and cell viability were normalized to the intensity of solvent control (DMSO) to obtain a percentage. The number of standard deviations far from the sample mean was obtained by z-standardization as follows: 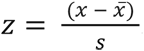, where z is the number of standard deviations far from the sample mean, *x* is the realization of fluorescence intensity, 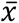 is the sample mean, and *s* is the standard deviation. The hierarchical clustering was performed in R (R version 4.0.2) with packages gplot using the normalized fluorescence and viability intensity as the clustering variable. For IC50 and EC 50 estimation, the protein expression was normalized to the expression of solvent control (DMSO) to obtain a percentage. The percentage against the log-transformation concentration was fitted to a four parameters logistic function in GraphPad Prism v7, and the value was calculated by the fitted model. For protein half-life and the degradation kinetics estimation, the protein expression was normalized to the expression of the initial time point to obtain a percentage. The percentage against the time was fitted to nonlinear regression in R with losses function in the stats package, and the half-life was calculated by the fitted model. Figures were generated in GraphPad Prism or ggplot2 package in R. Experiments were performed at least three times to obtain the statistical significance. The significant level was determined by the unpaired two-tail Student’s t-test in R. Data are presented as the mean ± sample error of the mean (SEM). P values < 0.05 were considered as significant.

## Acknowledgments

We thank Dr. Li-Rung Huang (National Health Research Institutes, Taiwan) for providing Ad-HBV, Dr. Yen, Hsueh-Chi Sherry (Institute of Molecular Biology, Academia Sinica, Taiwan) for providing protein stabiliy construct.

## Abbreviation

PreS2-LHBS: HBV large surface antigen with PreS2 region deletion
LHBS: large surface antigen
MHBS: middle surface antigen
SHBS: small surface antigen
CHB: Chronic hepatitis B
GGH: ground-glass hepatocyte
NA: nucleotide analog
NeHep-S2: NeHepLxHT stably expressing SNAP-tagged PreS2-LHBS
AdHBV: Adenoviral HBV gene transfer
EC_50_: 50% effective concentration
R_max_: maximal reduction percentage
GPS-PreS2-LHBS: 293T cells stably expressing GPS reported carrying PreS2-LHBS.

## Supporting information

**Figure S1.**
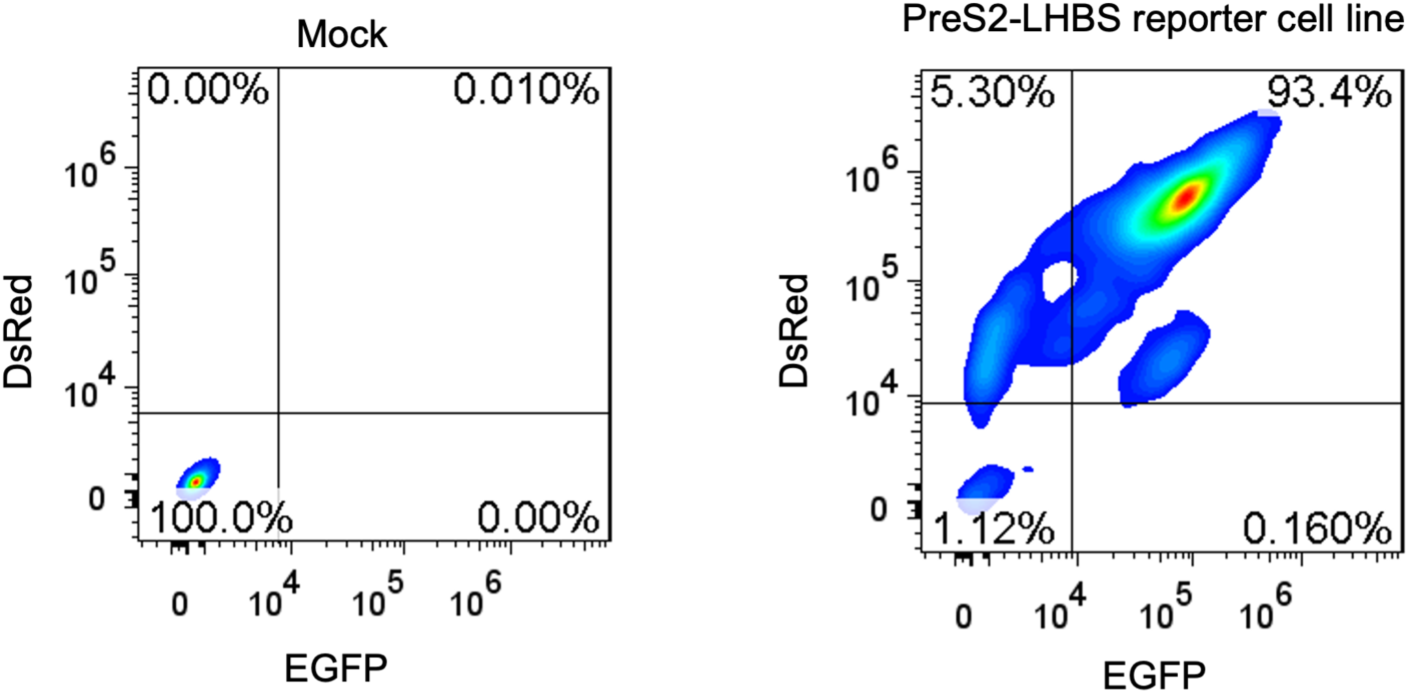
Expression profile of PreS2-LHBS reporter stable cell line. Ten thousand of 293T cells stably expressing PreS2-LHBS reporter were analyzed by flow cytometry. The population percentage located in each block was shown in the corner.

**Figure S2.**
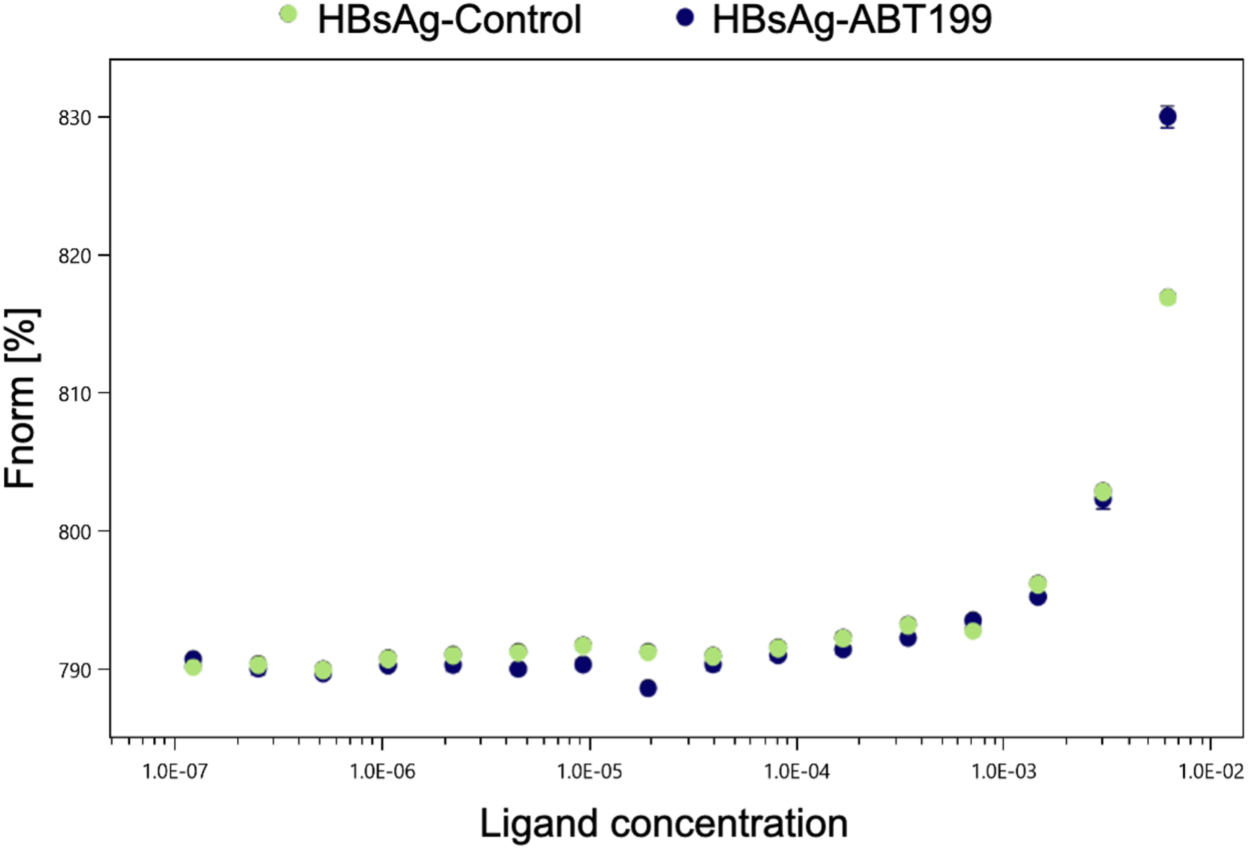
Investigation of the interaction between HBsAg and ABT199. 500nM of HBsAg protein were labeled with HisDye and were incubated with the indicated concentration of ABT199 and solvent control (DMSO). The graph shows the interaction signal determined by MST.

**Figure S3.**
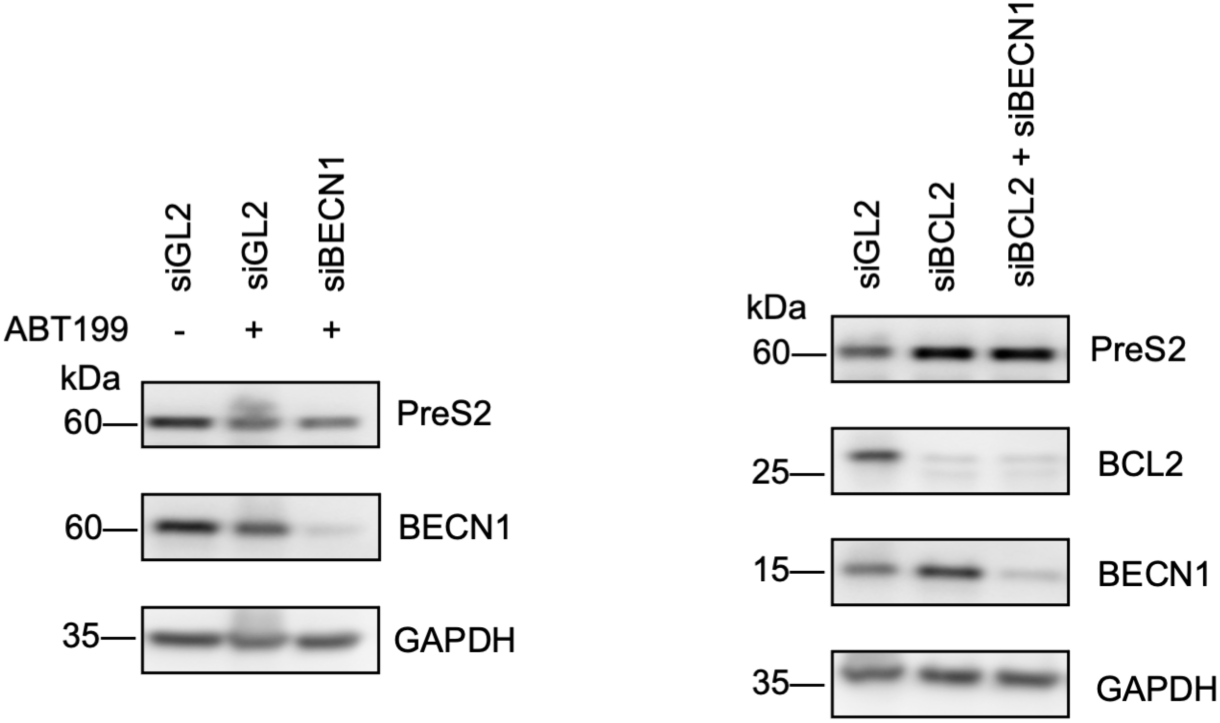
The effect of depleting BCL2 and BECN1. NeHep-S2 cells were knockdown by transfection with a BECN, BCL2, or control specific siRNA. The protein expression was then analyzed by immunoblotting. GAPDH served as a loading control.

**Figure S4.**
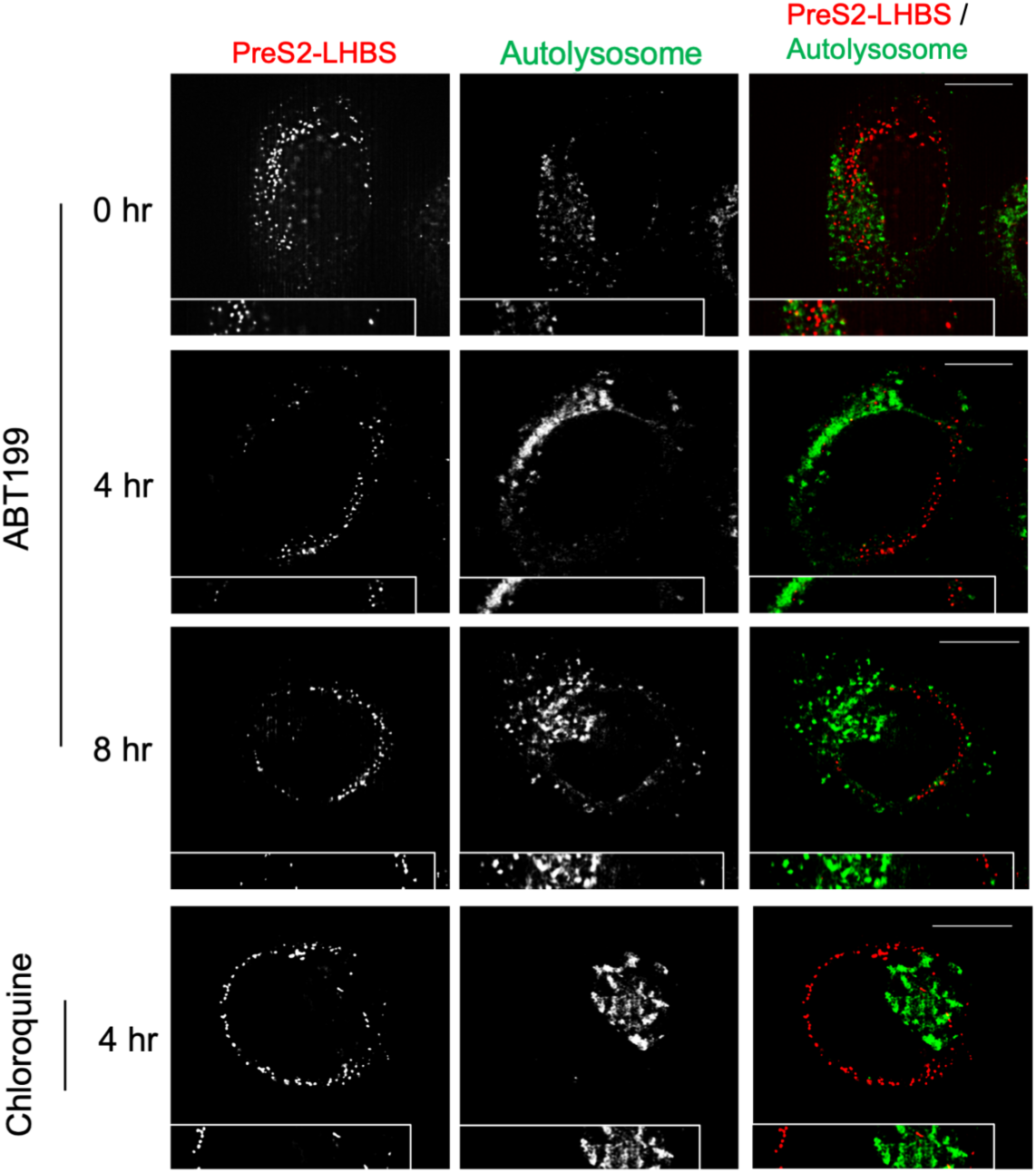
localization of macroautophagy and PreS2-LHBS upon ABT199 treatment. NeHep-S2 cells were stained for PreS2-LHBS and autolysosome after treating with ABT199 for indicated time points. Treatment of chloroquine served as a negative control. Scale bar donates 10um.

**Figure S5.**
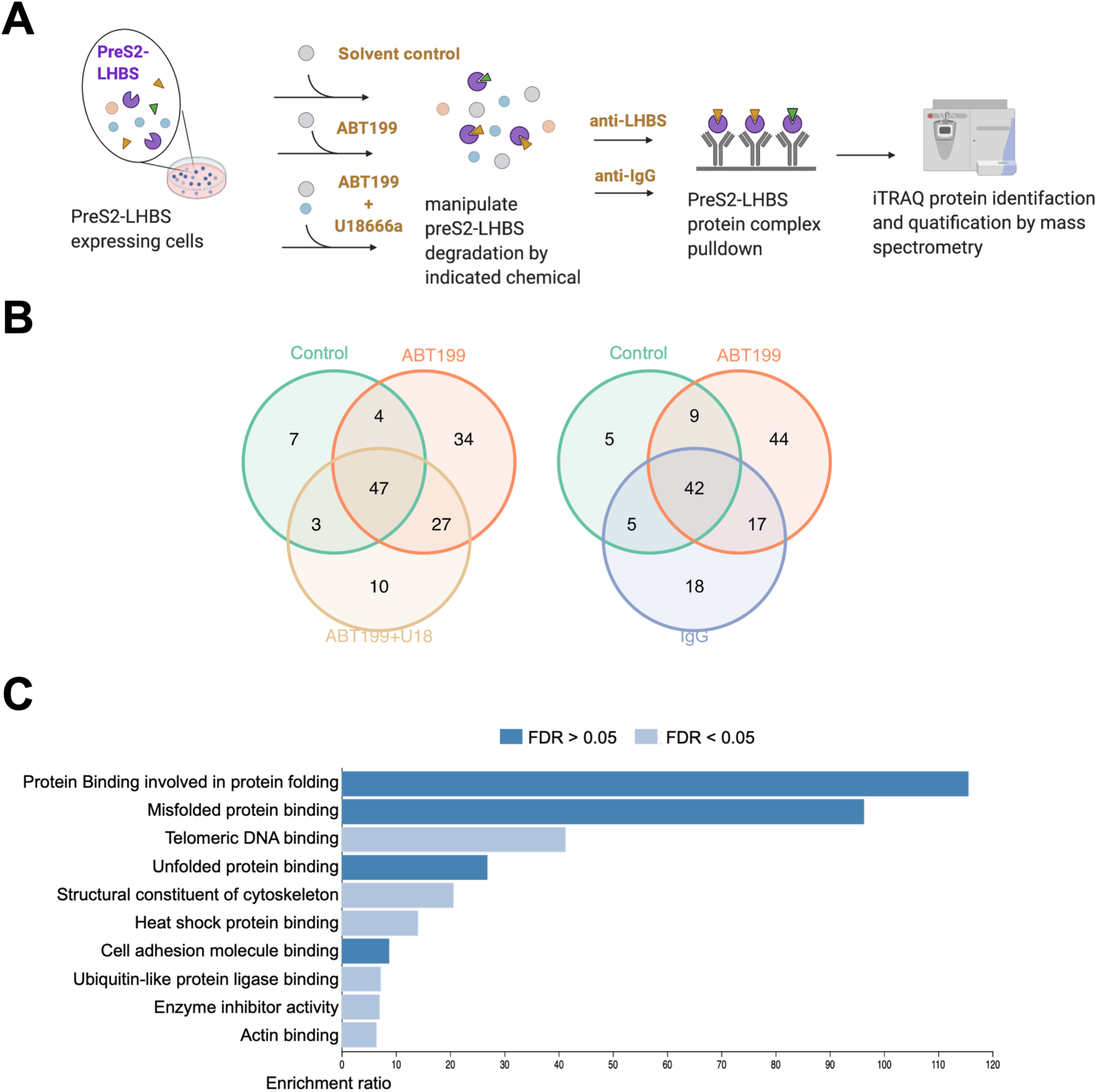
Purification screening of PreS2-LHBS interacted protein upon ABT199 treatment. (A) Schematic overview of FDA-approved drug library screening. (B) Venn diagram illustrates the relationship of identified protein in four treatments. ABT199 showed a significantly increased number of different proteins. (C) Functional enrichment analysis of the different proteins in ABT199 treatment.

**Table S1. FDA-approved drug screening result, related to Figure 1**

**Table S2. Identified protein in purification screening upon ABT199 treatment, related to Figure S5**

## Notes

### Competing Interest Statement

The authors have declared no competing interest.

